# A class of secreted mammalian peptides with potential to expand cell-cell communication

**DOI:** 10.1101/2023.06.02.543503

**Authors:** Amanda L. Wiggenhorn, Hind Z. Abuzaid, Laetitia Coassolo, Veronica L. Li, Julia T. Tanzo, Wei Wei, Xuchao Lyu, Katrin J. Svensson, Jonathan Z. Long

## Abstract

Peptide hormones and neuropeptides are fundamental signaling molecules that control diverse aspects of mammalian homeostasis and physiology. Here we demonstrate the endogenous presence of a sequence diverse class of orphan, blood-borne peptides that we call “capped peptides.” Capped peptides are fragments of secreted proteins and defined by the presence of two post-translational modifications – N-terminal pyroglutamylation and C-terminal amidation – which function as chemical “caps” of the intervening sequence. Capped peptides share many regulatory characteristics in common with that of other signaling peptides, including dynamic regulation in blood plasma by diverse environmental and physiologic stimuli. One capped peptide, CAP-TAC1, is a tachykinin neuropeptide-like molecule and a nanomolar agonist of multiple mammalian tachykinin receptors. A second capped peptide, CAP-GDF15, is a 12-mer peptide that reduces food intake and body weight. Capped peptides therefore define a largely unexplored class of circulating molecules with potential to regulate cell-cell communication in mammalian physiology.

## Introduction

Peptide hormones and neuropeptides are fundamental signaling molecules that mediate cell-cell communication (Hook et al., 2018). These signaling molecules are produced by proteolytic cleavage of secreted preproproteins and released extracellularly via the classical secretory pathway. Once secreted, peptide hormones and neuropeptides act on cognate receptors to regulate nearly all aspects of homeostasis and physiology. Because of their potent and powerful physiologic actions, peptide hormones and neuropeptides have attracted considerable pharmaceutical interest as starting points for the development of therapeutics across multiple human disease areas (Sanyal et al., 2018; Sikich et al., 2021; Wilding et al., 2021).

The biosynthesis of signaling peptides often involves post-translational enzymatic modification of C-and/or N-termini. Over 80 mammalian peptide hormones and neuropeptides possess a C-terminal amide instead of a free carboxylate (e.g., GLP-1) (Eipper et al., 1992; Uniprot Consortium, 2023). This modification is installed by the action of PAM (peptidyl-glycine alpha-amidating monooxygenase), an enzyme that cleaves off a C-terminal glycine to leave an amide terminus (Czyzyk et al., 2005; Eipper et al., 1983). In addition, chemical modifications of the N-terminus include N-acetylation (Guo et al., 2004), N-formylation (Chen et al., 2022), and N-pyroglutamylation (Lindemans et al., 2010). Combinations of C-and N-terminal modifications can also be found within the same peptide (Lindemans et al., 2010). In the cases where they have been examined, C-and N-termini modifications can both increase peptide stability and also enhance receptor affinity (Van Coillie et al., 1998; Goren and Bauce, 1977).

Historically, peptide hormone and neuropeptide-like molecules have been discovered via laborious and time-consuming activity-guided fractionation strategies (Harris and Lerner, 1957; Schally et al., 1971). More recently, shotgun mass spectrometry-based approaches using available mammalian genome information have been developed for the detection of peptide fragments from tissues and fluids (Fälth et al., 2007; Gupta et al., 2010; Tagore et al., 2009). However, the existing mass spectrometry methods typically identify few/most abundant peptides (e.g., low coverage), and consider only a selection of post translational modifications. In addition, the potential bioactivity and/or signaling of the peptides detected remains undefined.

We reasoned that knowledge of primary sequences indicative of post-translational processing of peptide termini, combined with a targeted mass spectrometry approach using authentic peptide standards, might enable de novo prediction and high-sensitivity detection of previously unknown peptide cleavage products. Importantly, these predicted cleavage products would harbor C-and N-terminal modifications that resemble that of other peptide hormones and neuropeptides, thereby suggesting also potential signaling activity. Using this strategy, here we report the discovery of a sequence-diverse class of orphan blood-borne mammalian peptides called “capped peptides.” Capped peptides are defined by the co-incident presence of two post-translational modifications, N-terminal proglutamylation and C-terminal amidation, that function as terminal “caps” of the intervening peptide sequence. Capped peptides also exhibit regulatory characteristics similar to other signaling peptides, including dynamic circulating levels in response to physiologic and environmental state. *In vitro* and *in vivo* functional assays establish signaling and bioactivity for two orphan capped peptides: CAP-TAC1 is a tachykinin neuropeptide-like molecule that exhibits nanomolar agonist activity at multiple mammalian tachykinin receptors, and CAP-GDF15, derived from the prepropeptide region of the anorexigenic hormone GDF15, is itself a novel 12-mer anorexigenic peptide. Our studies demonstrate that N-and C-terminal capping chemical motif that is present in a much larger number of endogenous secreted peptides than previously anticipated. Capped peptides therefore constitute a class secreted peptides with potential to broadly regulate cell-cell communication in mammalian physiology.

## Results

### Prediction and detection of capped peptides in mouse plasma

To define potential modified fragments from existing classically secreted proteins, we first used Uniprot (Consortium, 2023) to curate a collection of protein sequences corresponding to classically secreted mouse proteins (**Methods**). Our initial collection of N=2,835 sequences contained many known classically secreted proteins, including apolipoproteins, secreted enzymes, and preprohormones (**Table S1**). Next, we sought to define potential fragments of these sequences that might also harbor additional post-translational modifications of the N-and C-termini. For these modifications, we focused on N-terminal pyroglutamylation and C-terminal amidation (**Fig. 1A**). C-terminal amidation sequences were identified by the presence of a glycine-dibasic GRR/GKR tripeptide (**Fig. 1A**). N-terminal pyroglutamylation sequences were identified by the presence of a glutamine (Q) 3 to 20 amino acids upstream of the glycine-dibasic motif (**Fig. 1A**, **Table S2**).

**Fig. 1.**
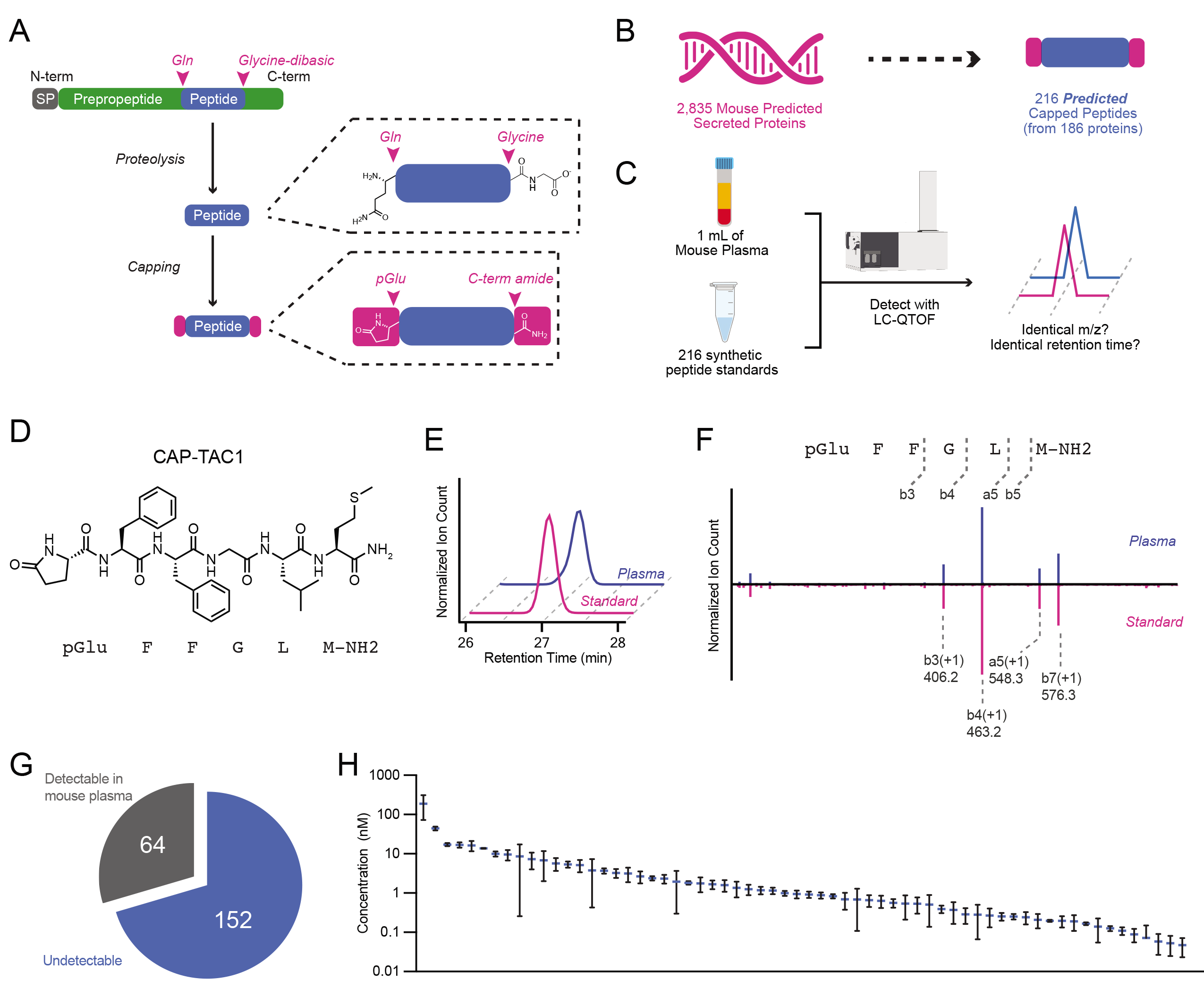
Genomic prediction and mass spectrometry detection of capped peptides in mouse plasma. (A) Schematic representation of capped peptide production from secreted, full-length preproprecursor proteins. N-term, N-terminal; C-term, C-terminal; SP, signal peptide; Gln, glutamine; pGlu, pyroglutamyl. (B,C) Number of predicted mouse capped peptides (B) and the targeted mass spectrometry strategy for their detection (C). (D) Chemical structure of CAP-TAC1 (pGlu-FFGLM-NH2). (E,F) Representative extracted ion chromatogram at m/z = 724.3 (E) and mirror fragmentation spectra (F) of authentic CAP-TAC1 standard (pink) and the endogenous mouse plasma peak (blue). (G, H) Pie chart showing number of detectable capped peptides in mouse plasma (G) and their quantitation (H). For (H), data are shown as means ± SEM (N=3 mice).

Our rationale for this set of prediction criteria, and for the focus on peptide fragments with coincident N-pyroglutamyl and C-amidation modifications, was based on the conceptual basis that: (1) these modifications have previously been shown to enhance receptor binding and plasma stability for other peptide hormones, therefore providing higher likelihood that our predicted sequences would be detectable by mass spectrometry and also exhibit bona fide signaling and bioactivity; (2) the glycine dibasic sequence is a well-established C-terminal cleavage motif that, after proteolysis, results in a C-terminal glycine which can subsequently be enzymatically modified via PAM to generate the C-terminal amide; (3) biosynthesis of other N-terminal pyroglutamylated signaling peptides often occurs via an N-terminal glutamine, rather than a glutamate. Using this computational framework, we predicted a total of 216 potential cleaved and modified mouse peptides from 186 classically secreted proteins encoded in the mouse genome (**Fig. 1B**). We call this class of modified peptides “capped peptides” owing to their terminal modifications which function as “caps” of the intervening peptide sequence (**Fig. 1A,B**). Our prediction of hundreds of such modified sequences suggests that chemical capping, rather than being unusual and rare modifications that have been previously reported in only three mammalian peptide hormones (GnRH, TRH, and gastrin), might in fact define a distinct chemical class and potentially functional motif present in signaling peptides.

To determine whether capped peptides are produced endogenously, we used a targeted mass spectrometry approach to directly measure the levels of all predicted capped peptides in mouse plasma. Our targeted mass spectrometry strategy was inspired by standard approaches in small molecule metabolomics, where analytes are confidently identified by comparison of retention times and mass-to-charge (m/z) ratios to authentic standards that were produced via chemical synthesis. Importantly, this mass spectrometry-based workflow obviates the need for identification via database searching that is classically associated with untargeted peptidomics or shotgun proteomics. Towards this end, we first isolated total plasma peptidome from mouse plasma. Briefly, mouse plasma was boiled to inactivate proteases, reduced with DTT, alkylated with iodoacetamide, and concentrated using a C8 column (see **Methods** for details). In parallel, we used solid phase peptide synthesis to generate authentic peptide standards for all 216 predicted capped peptides (**Fig. 1C**). A mixture of the 216 peptide standards was also reduced, alkylated, and concentrated. Total mouse plasma peptides were then compared to the synthetic peptide mixture by high resolution liquid chromatography-mass spectrometry (LC-MS) on a quadrupole-time-of-flight (qTOF) mass spectrometer (see **Methods** for details).

A representative example of a positive detection event for CAP-TAC1 (pGlu-FFGLM-NH2, **Fig. 1D**), a capped peptide derived from amino acids 63-68 of full-length TAC1 (protachykinin-1), is shown in **Fig. 1E**. Here, the authentic CAP-TAC1 standard exhibits identical m/z ratio and retention time to an endogenous plasma peak (m/z = 724.3, retention time = 27 min, **Fig. 1E**). As further validation of this structural assignment and detection event, MS/MS studies of the authentic CAP-TAC1 standard and the endogenous peak revealed nearly identical fragmentation patterns (**Fig. 1F**). Manual inspection of the daughter ions revealed that many could be directly assigned to the CAP-TAC1 sequence, including b3(+1, m/z = 406.2), b4(+1, m/z = 463.2), and a5(+1, m/z = 548.3) (**Fig. 1F**). Lastly, the endogenous m/z = 724.3 peak and our authentic standard of CAP-TAC1 exhibited a distinct retention time from that of an authentic standard of non-amidated CAP-TAC1 (pGlu-FFGLM-COOH, retention time = 28 min), further establishing our bona fide detection of the capped peptide (**Fig. S1**).

Across all predicted capped peptides, 64 of the 216 (30%) exhibited high confidence detection in blood plasma by mass spectrometry (**Fig. 1G** and **Table S2**). The remaining 152 capped peptides (70%) did not exhibit a peak at the retention time of the authentic standard. By comparison to an external standard curve, the detected mouse capped peptides exhibited circulating concentrations in the range of ∼0.1-100 nM (**Fig. 1H**). We conclude that capped peptides are a family of endogenously circulating molecules, much larger than previously described. Our inability to detect 152 of the predicted capped peptides may either reflect true absence of post-translational processing to generate those fragments, or alternatively, the endogenous presence of the capped peptide is at a level below our detection limit. Regardless, that many predicted capped peptides could in fact be endogenously detected by a targeted mass spectrometry pipeline also suggest that specific proteolytic processing and capping to produce protected peptide fragments is much more prevalent than previously anticipated.

### Sequence and gene-level analysis of mouse capped peptides

We next examined the sequences and set of full-length preproprecursor proteins of the detected capped peptides. By performing BLASTP alignment of the full capped peptide sequences with C-terminal GRR/GKR extensions, we could confirm that the vast majority of the detected mouse capped peptides detected (60/64, ∼94%) mapped uniquely to only one pre-proprecursor polypeptide sequence (**Fig. S2**). Two of the detected capped peptides, CAP-GNRH1 (pGlu-HWSYGLRPG-NH2) and CAP-GAST (pGlu-RPRMEEEEEAYGWMDF-NH2) directly corresponded with the known hormone sequences for GnRH and gastrin (**Fig. 2A, B**). These data demonstrate that our hybrid computational-analytical approach can “re-discover” two of the known signaling peptides that harbor both N-terminal pyroglutamylation and C-terminal amidation. An additional 23 detected capped peptides mapped to preproprecursor proteins and corresponding genes that had previously been annotated to generate polypeptides with signaling bioactivity, but for which a shorter cleavage fragment had not been previously identified. For instance, we detected a capped pentapeptide, CAP-FGF5 (pGlu-WSPS-NH2), derived from amino acids 77- 81 of the prepro-FGF5 (fibroblast growth factor 5) (**Fig. 2B**). A second capped peptide, CAP-GDNF (pGlu-AAAASPENSRGK-NH2) which mapped to amino acids 94-106 of GDNF (glial cell line-derived neurotrophic factor) (**Fig. 2B**). Classically, FGF5 is a so-called “paracrine” member of the fibroblast growth factor (FGF) family with little known about its function besides its involvement in hair follicle growth (Hébert et al., 1994). GDNF is a major growth factor that promotes the survival of dopaminergic and motor neurons; outside of the nervous system, GDNF is also a morphogen in the kidney and a spermatogonia differentiation factor (Airaksinen and Saarma, 2002). Our detection of CAP-FGF5 and CAP-GDNF suggests that FGF5 and GDNF might also exhibit endocrine functions via cleavage fragments generated the pre-proprecursor sequences. The remaining capped peptides (39/64, ∼61%) mapped to preproprecursors for proteins and genes that had not been previously suggested to have any roles in signaling. These include CAP-PLA2G2A (pGlu-FGEMIRLKT-NH2) which is cleaved from a phospholipase sequence and CAP-SNED1 (pGlu-STEVDRSVDRLTFGDLLP-NH2) which is derived from the poorly studied protein SNED1 (**Fig. 2B**).

**Fig. 2.**
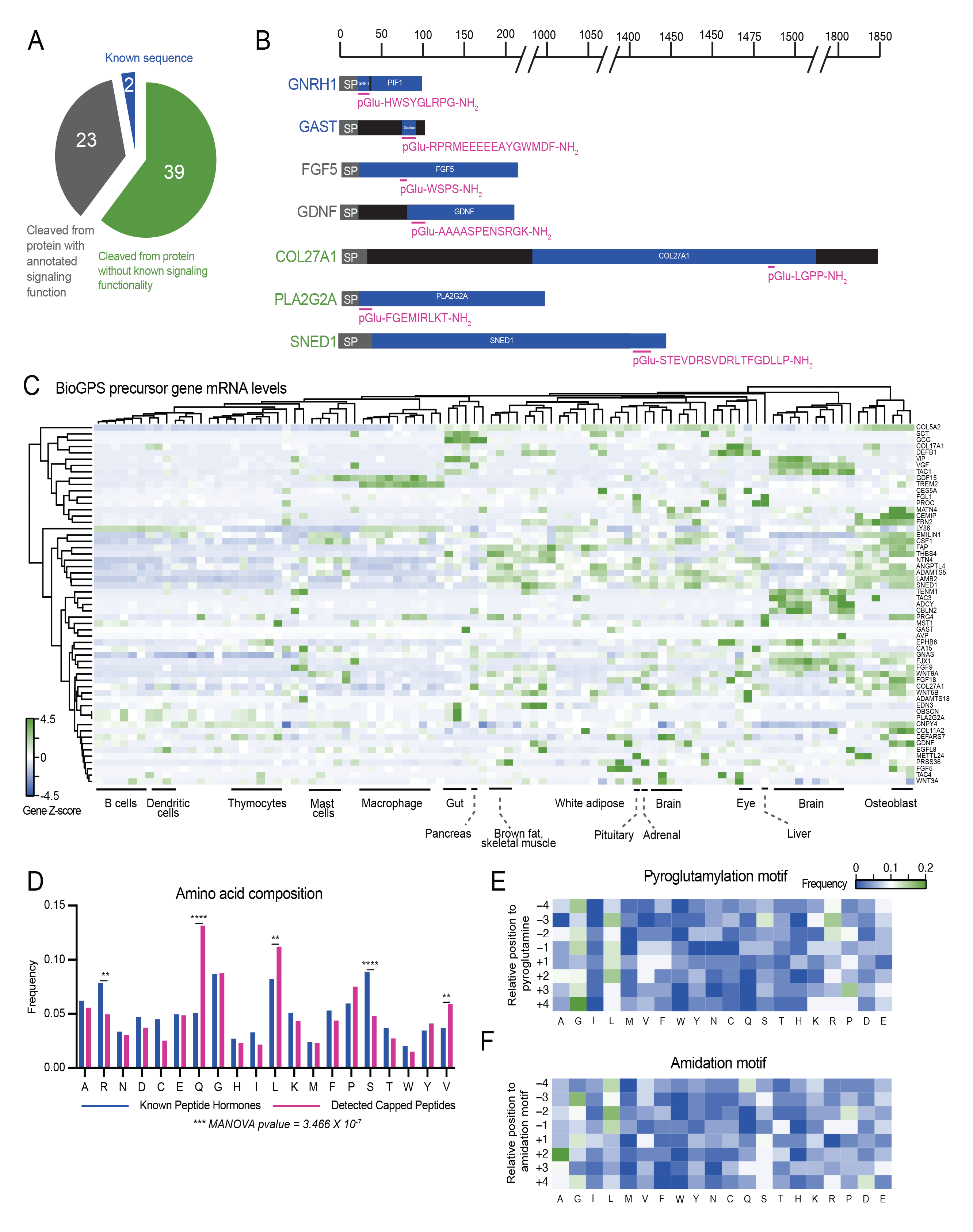
Sequence and gene-level analysis of capped peptides. (A,B) Pie chart (A) and representative examples (B) of the distribution of mouse capped peptides based on their full-length preproprotein precursors. SP, signal peptide. (C) H-clustered heat map of mRNA expression for capped peptide preproprecursor home genes across mouse tissues and cell types. (D) Frequency of each amino acid in capped peptides versus a Uniprot reference set of known peptide hormones and neuropeptides. (E, F) Heat map of amino acid frequency within four residues upstream and downstream of the N-terminal pyroglutamylation (E) or C-terminal amidation (F). For (C), data were obtained from BioGPS and shown as Z-score of the log-transformed value.

To further understand the cellular and tissue origin of capped peptides, we examined the mRNAs of the home genes encoding the capped peptide pre-proprecursors. We used BioGPS (Wu et al., 2016) as a reference mouse tissue gene expression dataset. As shown in **Fig. 2C**, this set of mRNAs exhibited both cell type-specific as well as widespread tissue expression. For instance, a strong enrichment of home gene mRNAs for certain capped peptides was found in the brain (e.g., CAP-CBLN2, CAP-TENM1, CAP-TAC3, CAP-ADCYAP1), in bone (e.g., CAP-MATN4, CAP-CEMIP), and in macrophages (e.g., CAP-GDF15, CAP-TREM2). Conversely, mRNAs corresponding to other capped peptides exhibited more diffuse tissue expression across multiple cell types and organs (e.g., CAP-VIP enrichment in both brain and gut and CAP-COL5A2 expression in > 10 tissues).

Lastly, we performed more detailed amino acid composition and sequence analysis of the detectable capped peptides from mouse plasma. As a reference and comparison set, we once again used Uniprot to manually curate a set of known mouse peptide hormones and neuropeptides (see **Methods** and **Table S3**). Glutamine was enriched in capped peptides compared to the reference set of known peptide hormones and neuropeptides, which was expected based on our original computational search criteria. In addition, two hydrophobic amino acids, valine and leucine, were also more prevalent in capped peptides, whereas two polar amino acids (arginine, serine) were less represented (**Fig. 2D**). To understand whether there might be additional sequence-specific determinants of capping beyond our original N-terminal Q and C-terminal GRR/GKR motifs, we examined the amino acid sequences centered around the N-and C-termini. A modest enrichment of glycine and leucine were observed to flank both the N-terminal pyroglutamylation motif (**Fig. 2E**), differing from the dibasic residue cleavages sites typically flanking hormones. In addition to glycine and leucine enriched around the C-terminal amidation motif, we also observed a strong enrichment for alanine at the +2 position (**Fig. 2F**). Together, these expression and sequence analyses demonstrate that capped peptides are likely produced from diverse tissues and exhibit specific patterns of amino acid composition and sequence.

### Dynamic regulation of capped plasma levels in mice

Many signaling peptides exhibit dynamic regulation in a manner dependent on internal physiologic state or external environmental conditions. We therefore measured the circulating levels of capped peptides after six distinct perturbations that spanned a wide range of physiologic processes, environmental stimuli, organ systems, and time scales: 16 h fasting vs. fed, 8-weeks high fat diet feeding vs. chow feeding, lipopolysaccharide (LPS, 0.5 mg/kg, intraperitoneal) vs. vehicle, 6AM vs. 6PM, acute treadmill running (1 h) vs. sedentary, and 3 months vs. 24 months old. For each comparison, mouse plasma was collected and processed as described previously, and capped peptides were quantified by LC-MS (**Fig. 1**).

As shown in **Fig. 3A**, each physiologic comparison resulted in bidirectional regulation of a unique subset of capped peptides. Across all measurements (N=384, corresponding to N=64 capped peptides in each of N=6 conditions), we observed a total of N=93 (24%) instances for a particular capped peptide being elevated (N=39) or suppressed (N=54) by more than 2-fold in any condition. 73% (47/64) of the capped peptides exhibited at least one instance of 2-fold change across any condition; similarly, 100% of the conditions (6/6) yielded at least one capped peptide that was changed by more than 2-fold.

**Fig. 3.**
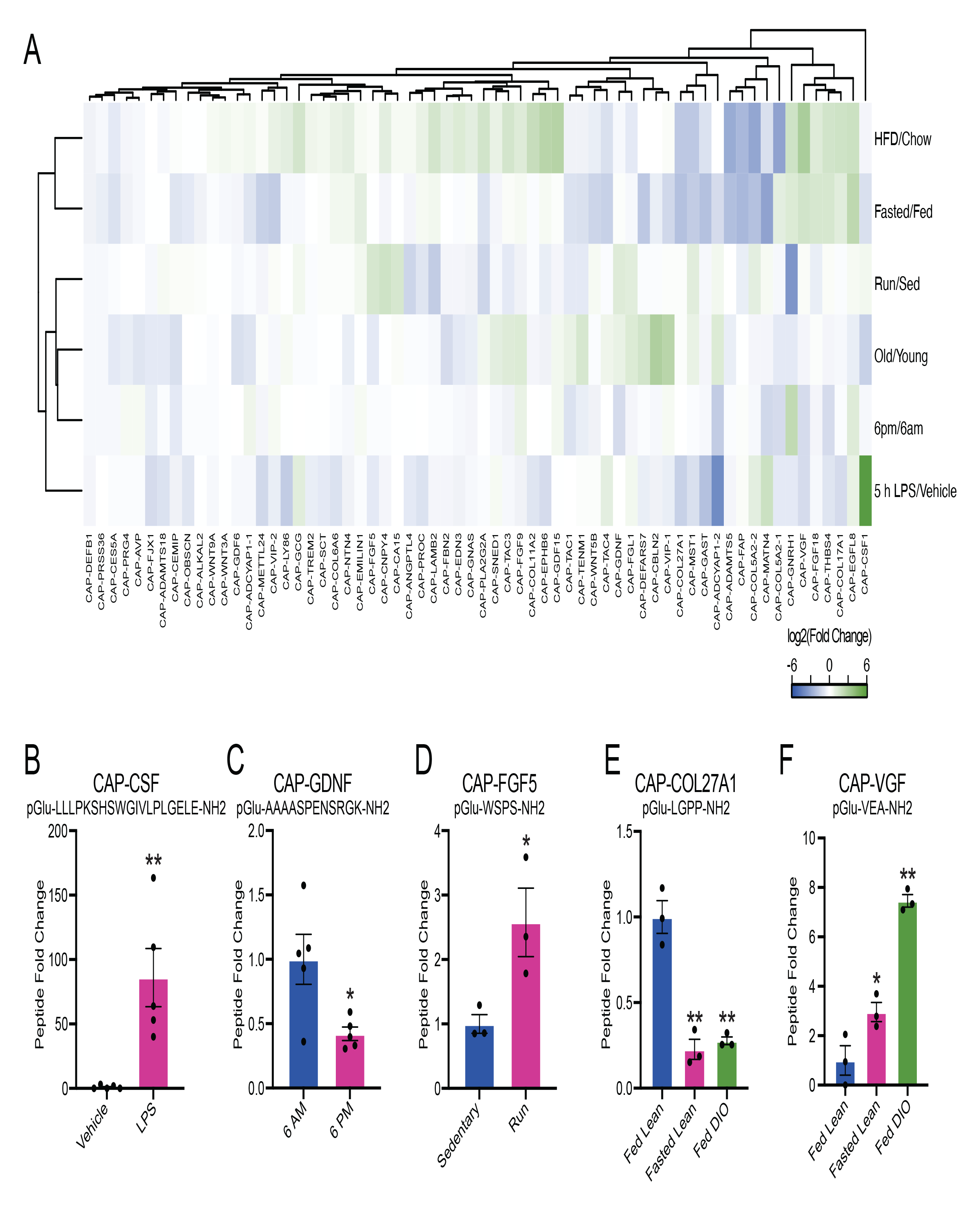
Dynamic regulation of circulating capped peptide levels by different environmental and physiologic perturbations. (A) Heat map of the relative fold change for the indicated capped peptide in the indicated condition. (B-F) Representative peptide quantification for the indicated capped peptide in the indicated comparison. For (A-F), “Fasting/Fed” indicates a comparison between plasma from 3- month old male mice on chow diet versus age-and sex-matched controls after fasting for 16 h (N=3/group); “HFD/Chow” indicates a comparison between plasma from 4-month old male mice on chow diet versus 4-month old male mice after being on high fat diet (HFD, 60% kcal from fat) for 9 weeks (N=3/group); “LPS” indicates a comparison between plasma from 2-month old male mice following treatment with vehicle versus LPS (0.5 mg/kg, IP, 5 h, N=5/group); “6AM/6PM” indicates a comparison between plasma from 2-month old male mice at the beginning of the night cycle (6AM) versus age-and sex-matched mice at the beginning of the day cycle (6PM) (N=5/group); “Old/Young” indicates a comparison between plasma from 4-month old male mice versus 23-month old male mice (N=2/group); “Run/Sed” indicates a comparison between plasma from sedentary 3-month old male mice versus age-and sex-matched controls after an acute exhaustive 60 min run on a treadmill. For (B-F), data are shown as means ± SEM. * P < 0.05, ** P < 0.01 versus control by Student’s t-test.

The capped peptide/perturbation pair resulting in the most dramatic regulation was CAP-CSF1 (pGlu-LLLPKSHSWGIVLPLGELE-NH2), derived from amino acids 419-438 of full-length prepro-CSF1. Plasma CAP-CSF1 levels were induced by ∼84-fold after LPS treatment (P < 0.01, **Fig. 3A, B**). Importantly, CAP-CSF1 levels were unchanged in any of the other comparisons (**Fig. 3A**), establishing that induction of CAP-CSF1 in plasma is a specific response to an inflammatory stimulus. Previously, the most well-known polypeptide product derived from full-length prepro-CSF1 is m-CSF1 (macrophage colony-stimulating factor 1), which is itself an LPS-inducible cytokine (Benmerzoug et al., 2018). The co-induction of CAP-CSF1 may therefore represent additional, LPS-inducible proteolytic processing of m-CSF1. In addition, we could also identify several other interesting examples of individually regulated dynamic peptides in each of the conditions. For instance, CAP-GDNF was selectively downregulated in plasma collected at 6PM versus 6AM (pGlu-AAAASPENSRGK-NH2, 58% reduction, P < 0.05, **Fig. 3C**) and CAP-FGF5 was selectively induced by a single bout of treadmill running (1 h) versus sedentary mice (pGlu-WSPS-NH2, 2.6-fold increase, P < 0.05, **Fig. 3D**).

Beyond high magnitude changes in individual capped peptides in each condition, we also identified examples of capped peptides that exhibited coordinate regulation across multiple physiologic states. For instance, we observed a cluster capped peptides that were coordinately regulated in two distinct nutritional stressors, fasting and high fat diet feeding. Individual examples of dynamic capped peptides within this nutrition-regulated cluster included CAP-COL27A1 (pGlu-LGPP-NH2, ∼75% reduction, P < 0.05, **Fig. 3E**) and CAP-VGF (pGlu-VEA-NH2, >3-fold induction, P < 0.05, **Fig. 3F**). The nutritional regulation of this subset of capped peptides, and of CAP-COL27A1 and CAP-VGF in particular, might point to specific functions in nutrient harvesting, fuel metabolism, or energy homeostasis. Together, we conclude that capped peptide levels in the circulation are dynamically regulated in a manner dependent on the specific capped peptide and specific physiologic perturbation.

### CAP-TAC1 is a novel tachykinin with homology to Substance P

Our data so far suggest capped peptides exhibit many structural and regulatory features of other well-established peptide hormones and neuropeptides. We next sought to determine whether any of the detectable capped peptides exhibited signaling and/or functional bioactivity. We first focused on CAP-TAC1 (pGlu-FFGLM-NH2). As we already showed in **Fig. 1**, CAP-TAC1 is robustly detected in blood plasma. The full-length TAC1 preproprotein encodes multiple members of the tachykinin neuropeptides, including Neurokinin A/Substance K, Neuropeptide K/Neurokinin K, Neuropeptide gamma, and Substance P (**Fig. 4A**) (Steinhoff et al., 2014). Previously, C-terminal fragments of Substance P have been reported in the context of pharmacological studies. For instance, FGLM-NH2 (4-11) and QFFGLM-NH2 (6-11) are potent tachykinin receptor agonists (Krumins et al., 1993). In addition, a synthetic fragment C-terminal fragment pGlu-FFPLM-NH2 (e.g., SP[6-11,pGlu^6^,Pro^9^], “septide”) is a chemical agonist of the tachykinin recpetors that binds to a site distinct from full-length Substance P (Pradier et al., 1994; Hastrup et al., 1996; Bellucci et al., 2002). Other C-terminal fragments of Substance P have also been detected endogenously (e.g., SP[5-11]) (Sakurada et al., 1985). Notably, none of these prior studies have described the unique chemical structure of CAP-TAC1 (e.g., pGlu-FFGLM-NH2), which contains both pyroglutamyl and amidation caps around the C-terminal amino acids 6-11 of Substance P. Therefore CAP-TAC1 is a novel circulating polypeptide with homology to fragments of Substance P.

**Fig. 4.**
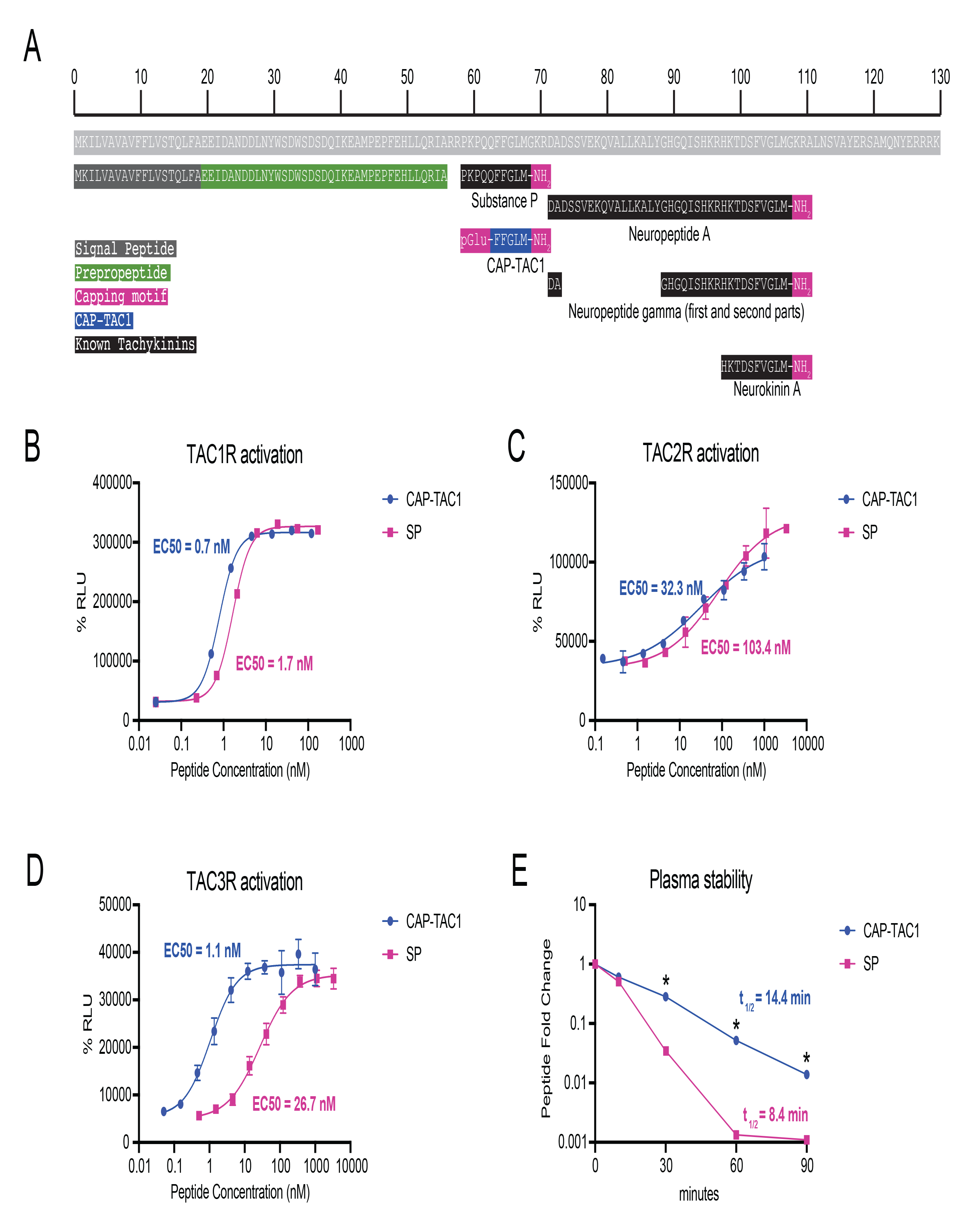
CAP-TAC1 is a potent tachykinin receptor agonist. (A) Schematic and annotation of the primary amino acid sequence for full-length mouse TAC1 preproprotein and its cleavage products. For (B-D), chemiluminescent signal intensity in PathHunter beta-arrestin CHO-K1 cells transfected with human (B) TACR1, (C) TACR2, and (D) TACR3 following treatment with the indicated concentration of CAP-TAC1 or substance P (SP). (E) Relative levels of CAP-TAC1 or substance P (SP) in mouse plasma following incubation at 37°C for the indicated time. For (B-D), N=2 per concentration; for (E), N=3 per condition. For (B-D), data are shown as means ± SEM. * P < 0.05, ** P < 0.01 versus control by Student’s t-test.

CAP-TAC1’s homology to substance P suggested that this novel peptide might also function as a tachykinin neuropeptide-like agonist. We therefore used a cellular human TACR1- beta arrestin recruitment assay with a fluorescence readout to directly determine the ability of CAP-TAC1 to agonize the TACR1 (also called NK1R), a high affinity receptor for substance P (Bhatia et al., 1998). As shown in **Fig. 4B**, CAP-TAC1 exhibited dose-dependent and high potency agonism of TACR1 (EC50 = 0.7 nM). As a positive control, substance P exhibited a similar dose-dependent activation (EC50 = 1.7 nM). Both CAP-TAC1 and substance P exhibited similar levels of maximal activation (CAP-TAC1, 96.6% of maximal response; substance P, 99.8% of maximal response).

In addition to TACR1, there are two other mammalian tachykinin receptors, TACR2 (NK2R) and TACR3 (NK3R). Remarkably, CAP-TAC1 was 3-fold and 27-fold more potent than that of the positive control Substance P (CAP-TAC1, EC50 = 32 nM and 1 nM for TACR2 and TACR3, respectively; Substance P, EC50 = 103 nM and 27 nM for TACR2 and TACR3, respectively, **Fig. 4C,D**). CAP-TAC1 also exhibited comparable, but slightly different maximal activation of these two receptors versus Substance P (CAP-TAC1, 70% and 114% of the maximal response for TACR2 and TACR3, respectively; Substance P, 100% and 98% of the maximal response for TACR2 and TACR3, respectively). We conclude that CAP-TAC1 is a full agonist of multiple mammalian tachykinin receptors with potency similar to, or in some cases higher than previously established tachykinin neuropeptides.

Beyond differences in TACR activation, we reasoned that the two chemical caps of CAP-TAC1 might also produce important functional differences in terms of stability and resistance to proteolytic degradation compared to Substance P. To directly test this possibility, CAP-TAC1 (10 µM) and substance P (10 µM) were individually incubated with mouse plasma and incubated at 37°C. and their levels over time were measured by LC-MS. Substance P exhibited time-dependent degradation with a *t*_1/2_ = 8.4 min. By contrast, the rate of CAP-TAC1 degradation was substantially slower (*t*_1/2_ = 14.4 min) (**Fig. 4E**). In fact, levels of CAP-TAC1 were still detectable after 90 min, a time point when substance P was undetectable (**Fig. 4E**). Together, these data demonstrate that CAP-TAC1 exhibits similarities (e.g., tachykinin receptor agonism) as well as important differences (e.g., increased potency, increased plasma stability) compared to previously described tachykinin neuropeptides.

### A 12-mer anorexigenic capped peptide from the preproprecursor region of GDF15

We next sought to understand whether signaling and bioactivity might be indeed a general feature of many capped peptides beyond CAP-TAC1 alone by performing functional studies of a second orphan capped peptide, CAP-GDF15 (pGlu-LELRLRVAAGR-NH2, **Fig. 5A**). Full-length GDF15 is a secreted, 303 amino acid preproprecursor that, upon cleavage at R188, produces a C-terminal 114-amino acid anorexigenic protein hormone, which is also called GDF15 (Chrysovergis et al., 2014; Johnen et al., 2007; Macia et al., 2012). Interestingly, CAP-GDF15 mapped to amino acids 174-185, a region just upstream of the canonical GDF15 hormone and localized in the GDF15 prepropeptide region (**Fig. 5A**). CAP-GDF15 eluted at the same retention time as an authentic standard (**Fig. 5B**). In addition, tandem mass spectrometry fragmentation also revealed a similar fragmentation pattern between the endogenous peak and the authentic standard (**Fig. 5C**). These data demonstrate that full-length GDF15 precursor in fact encodes at least two polypeptide products.

**Fig. 5.**
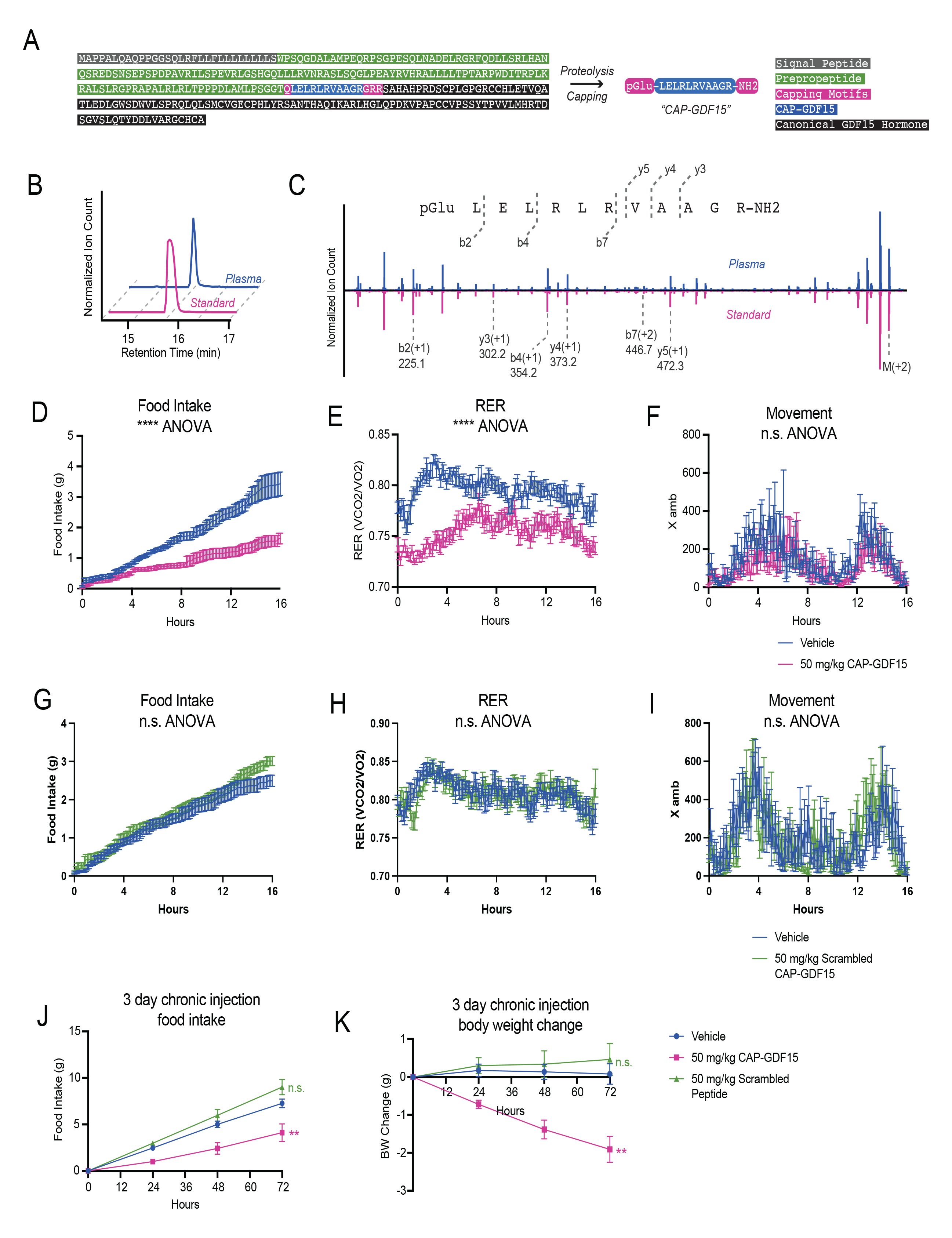
Functional studies of CAP-GDF15 in feeding and energy balance in mice. (A) Schematic of the primary amino acid sequence for full-length mouse GDF15 preproprotein and annotation of CAP-GDF15 sequence as well as the canonical GDF15 hormone sequence. (B, C) Representative extracted ion chromatogram (B) and mirror tandem fragmentation spectra (C) of CAP-GDF15 (pGlu-LELRLRVAAGR-NH2; +2 ion m/z = 682.4) and the endogenous plasma peak. (D-F) Food intake (D), respiratory exchange ratio (RER, E), and ambulatory activity (F) of 12-16-week diet-induced obese male mice following a single treatment of CAP-GDF15 (50 mg/kg, intraperitoneal) or vehicle control. (G-I) Food intake (G), RER (H), and ambulatory activity (I) of 12-16-week diet-induced obese male mice following a single treatment of scrambled CAP-GDF15 (50 mg/kg, intraperitoneal, scrambled sequence: pGlu-GLEALRARLRV-NH2) or vehicle control. (J, K) 3-day food intake (J) and body weight (K) of 12-16-week diet-induced obese male mice following treatment with CAP-GDF15 or scrambled CAP-GDF15 (50 mg/kg/day, IP), or vehicle control. Data are shown as means ± SEM. For (D-F), N=12/group; for (G-I), N=8/group; for (J,K), N=7/group. For (D-I), injection occurred at time T=0 (5:00pm) and data was collected for the following 16 hrs. For (D-K), ** P < 0.01, *** P < 0.001 by two-way ANOVA.

Unlike CAP-TAC1, the amino acid sequence of CAP-GDF15 did not immediately provide insights into its potential functions. However, we reasoned that the hypophagic effects previously demonstrated by overexpression of full-length GDF15 might extend beyond the classical GDF15 hormone alone to also include CAP-GDF15. To test this possibility, we administered a single dose of CAP-GDF15 (50 mg/kg, intraperitoneally) to diet-induced obese mice and measured whole body parameters of energy balance in metabolic chambers. CAP-GDF15 strongly suppressed food intake by ∼60% compared to vehicle-treated mice (**Fig. 5D**). A corresponding and expected suppression of respiratory exchange ratio (RER) was also observed (**Fig. 5E**). Notably, CAP-GDF15 did not alter movement (**Fig. 5F**), oxygen consumption (VO2, **Fig. S3**), or carbon dioxide production (VCO2, **Fig. S3**), demonstrating that the pharmacological effects of this peptide are specific to feeding control rather than other pathways of energy expenditure.

We next synthesized a control CAP-GDF15 peptide that preserved amino acid composition but scrambled the intervening amino acid sequence (scrambled CAP-GDF15, pGlu-GLEALRARLRV-NH2). This scrambled peptide control was completely ineffective in suppressing food intake and RER in metabolic chambers under identical experimental conditions (**Fig. 5G-I** and **Fig. S1**). We conclude that the anorexigenic effects of CAP-GDF15 are specific to this amino acid sequence.

Lastly, to determine whether the acute hypophagic effects of CAP-GDF15 would lead to long-term suppression of feeding and obesity, we administered CAP-GDF15 or scrambled CAP-GDF15 (50 mg/kg/day, IP), or vehicle control to diet-induced obese mice. Food intake and body weight were monitored over a three-day period. A durable suppression of food intake in CAP-GDF15-treated mice was over the three-day experiment (Vehicle, 7.3 ± 0.5 g/mouse; CAP-TAC1, 4.1 ± 0.9 g/mouse/day, P < 0.05, **Fig. 5J**). Consequently, and as expected, an increasing reduction in body weight was also detected (Vehicle, +0.1 ± 0.3 g/mouse; CAP-TAC1, −1.9 ± 0.3 g/mouse, P < 0.001, **Fig. 5K**). Importantly, mice treated with the scrambled CAP-GDF15 peptide were indistinguishable in body weight or food intake from control mice (P > 0.05 versus vehicle-treated mice, **Fig. 5J,K**). These data show that chronic CAP-GDF15 administration suppresses food intake and reduces body weight in a sequence-dependent manner. Together with CAP-TAC1, these data on CAP-GDF15 provide functional evidence for the signaling and bioactivity of two orphan capped peptides in both cell and animal models.

### Detection of human capped peptides and sequence comparison to mice

The capped peptide discovery pipeline described here only requires a full genome sequence and authentic peptide standards. Therefore, such an approach should also be readily amenable for discovering capped peptides in other species. Towards this end, we used the same hybrid computational-biochemical workflow as shown in **Fig. 1**, but now applied to protein sequences corresponding to classically secreted human proteins. Starting from N=3,791 secreted proteins, we predicted a total of 260 potential human capped peptides from 231 proteins (**Table S4**, **Table S5**, and **Fig. S4A**). We synthesized authentic peptide standards by solid phase peptide synthesis corresponding to all 260 possible human capped peptides (**Methods**). Once again, the retention time and parent masses of all authentic capped peptides were compared to endogenous peaks from commercially available pooled human plasma. In total, we robustly detected N=85 human capped peptides by mass spectrometry (**Fig. 6A**), a number that, by percentage, is similar to that previously observed with mice (30% detected/predicted for mouse, and 33% detected/predicted for human). Human capped peptides exhibited a similar distribution in plasma abundance (as quantitated using an external standard curve, **Fig. S4B**) and similar sequence characteristics in terms of amino acid composition (**Fig. S4C,D**) as mouse capped peptides. Using GTEx as a reference gene expression dataset, human capped peptides were also derived from preproprecursors whose mRNA levels also exhibited tissue-restricted, as well as more broad expression (**Fig. S5**).

**Fig. 6.**
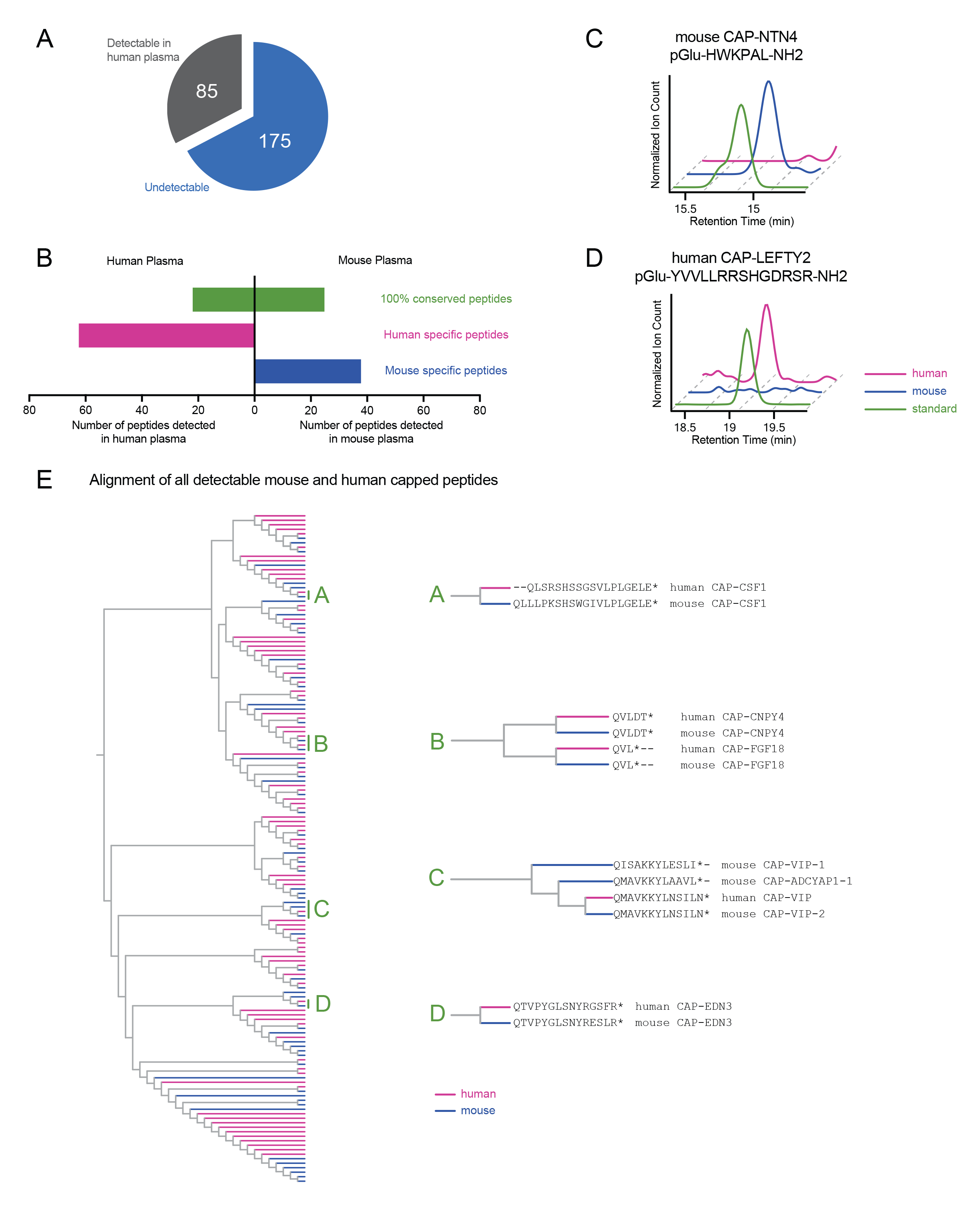
Detection of human capped peptides and sequence alignment comparison to mouse. (A) Pie graph of detectable human capped peptides. (B) Number of human or mouse capped peptides detected in human or mouse plasma, stratified by those capped peptide with 100% conservation (green) or those that are not 100% conserved (human-specific, purple; mouse-specific, blue). (C,D) Representative extracted ion chromatograms of the indicated capped peptides in plasma from the indicated species. (E) Phylogenetic alignment of all detectable capped peptides in human and mouse, with primary capped peptide sequences for select subclusters A-D shown on the right.

Because of sequence differences between the mouse and human proteome, we predicted that the set of humans capped peptides should be overlapping, but still distinct compared to those present in mouse plasma. Indeed, we detected many capped peptides in both human and mouse plasma that were 100% sequence conserved. In addition, none of the 64 human-specific peptides were detected in mouse plasma, and none of the 38 mouse-specific peptides were found in human plasma (**Fig. 6B**). Representative extracted ion chromatograms corresponding to conserved and species-specific capped peptides are shown in **Fig. 6C,D**.

Next, we performed a multiple sequence alignment to globally understand the sequence relationship and homology across all capped peptide sequences from both mice and humans. We also performed this analysis to understand whether the human-and mouse-specific capped peptide constituted entirely distinct sequences, high homologous sequences, or some combination of these two possibilities. The resulting dendrogram is shown in **Fig. 6E**. We selected several sub-clusters as illustrative examples here. Clusters “A” consists of a pair of capped peptides, mouse and human CAP-CSF1. These two are highly homologous sequences derived from amino acids 419-438 and 424-441 of full-length mouse and human CSF1, respectively. However, because the sequences are not 100% identical, mouse and human CAP-CSF1 are considered species-specific in our analysis. A similar example is shown in Cluster “D” where mouse and human CAP-EDN3 differ only by two amino acid residues and are once again considered species-specific, homologous sequences. Clusters “A” and “D” demonstrate that at least a subset of the species-specific sequences is due to differences in amino acid sequences of the corresponding full-length preproproteins from which the capped peptides are derived.

In another example, cluster “B” contained four short 3- and 5-mer capped peptides, which were amongst the shortest sequences in the entire dataset. The 3-mer capped peptides (pGlu-VL-NH2) were derived from the full-length mouse and human FGF18 sequences and exhibited identity between the two species. The other two capped peptides, derived from CNPY4 and again identical between mouse and human, constitute CAP-FGF18 homologs with an aspartyl-threonyl C-terminal extension (pGlu-VLDT-NH2). This cluster demonstrates that highly homolgous capped peptides can also be produced from distinct full-length preproprotein precursors.

Lastly, the cluster labeled “C” contained four peptides, three of which were derived from the full-length mouse or human VIP preproprecursor. The three VIP-derived capped peptides correspond with C-terminal fragments of the known PHI-27 and VIP peptide hormones. We also identified a non-VIP-derived peptide, CAP-ADCYAP1, which also exhibited high sequence alignment within this CAP-VIP-enriched cluster. CAP-ADCYAP1 is a C-terminal fragment of the neuropeptide PACAP (pituitary adenylate cyclase-activating polypeptide). These data suggest that the similar signal transduction pathways of VIP and PACAP peptides might also extend to additional fragments of these canonical sequences. We conclude that at least a subset of the human-and mouse-specific capped peptides represent highly homologous sequences. These data also globally identify similarities as well as important differences in the sequences of capped peptides between two species.

## Discussion

Here we provide multiple lines of evidence to show that capped peptides constitute a large class of previously unstudied mammalian signaling peptides. First, capped peptides are endogenously present in mouse and human plasma, where their levels are dynamically regulated by physiologic perturbations. Second, capped peptides exhibit post-translational N-and C-terminal modifications (pyroglutamylation and amidation, respectively) that resemble that of other peptide hormones and neuropeptides. Third, functional studies for two previously orphan capped peptides uncovered a tachykinin neuropeptide-like molecule as well as a novel anorexigenic peptide, demonstrating functional bioactivity for at least two members of this class. Lastly, the vast majority of the precise capped peptide sequences reported here have not been previously described as chemically defined, endogenous substances in mammals. In fact, our detection of >100 capped peptides in mouse and human plasma suggests that the N-and C-terminal “capping,” rather than being unusual and rare post-translational modifications, instead defines a distinct chemical motif that is present in a family of peptides with potential to mediate diverse axes of cell-cell communication.

Our ability to endogenously detect capped peptides was enabled by a custom mass spectrometry pipeline that uses a targeted mass spectrometry approach with authentic peptide standards. This method was inspired by classical approaches in targeted metabolomics, where small molecule mass-to-charge ratios and retention times are routinely compared against synthetic standards. The generality and simplicity of this approach was demonstrated by profiling the capped peptides present in blood plasma from two different species. Importantly, such a targeted approach obviates the need for large-scale database searching, thereby overcoming the problem of potential false discovery rate over-correction which may exclude true positive detection events. Because of potential limits of detection, it is not unreasonable to imagine that additional capped peptides that were undetectable here might indeed be endogenously present using more sensitive mass spectrometry methods. Projecting forward, these data also suggest that more targeted mass spectrometry approaches using synthetic standards, particularly on subsets of privileged peptide modifications and/or sequences, might be a powerful approach to identify peptides of specific structure and/or sequence within large, complex mass spectrometry datasets.

Functional studies of two orphan capped peptides, CAP-TAC1 and CAP-GDF15, provides new insights into cell-cell communication in two distinct areas of signaling and physiology. Nearly all tachykinin neuropeptides discovered to date have been identified by classical biochemical purification approaches. Our identification of CAP-TAC1, and then demonstration of this molecule as a high affinity agonist of mammalian tachykinin receptors, shows that additional fragments of full-length tachykinin preproproteins may be important endogenous mediators tachykinin signaling. Interestingly, CAP-TAC1 is only 5 amino acids in length and therefore represents the shortest naturally-occurring tachykinin receptor agonist reported to date in any species. Like CAP-TAC1, the detection of CAP-GDF15 also demonstrates that a single full-length preproprecursor (in this case, full-length GDF15) can generate more than a single bioactive polypeptide product. The observation that CAP-GDF15 is an anorexigenic peptide like the canonical GDF15 hormone raises new questions about the relative physiologic contribution of each CAP-GDF15 and canonical GDF15. In addition, because the sequences are largely distinct, we suspect that the downstream receptor(s) of CAP-GDF15 are likely to be distinct from that of the canonical GDF15 hormone.

Beyond these two specific examples, a major future challenge and goal will be to annotate the signaling and potential functions for other capped peptides. Already, our studies of CAP-TAC1 and CAP-GDF15 provide a potential roadmap for these studies. First, several other capped peptides are similar to CAP-TAC1 in that they represent smaller fragments of known peptide hormones. It is not unreasonable to imagine that these other capped peptides engage at the corresponding hormone and neuropeptide receptors as the canonical full-length signaling peptides. Second, potential functional hypotheses might arise from analysis of the physiologic functions of the full-length proteins, especially those that might not be yet explained via the action of the canonical proteins. Lastly, full-scale screening of capped peptides against a panel of candidate G-protein coupled receptors may also uncover the corresponding receptors for individual capped peptides. We anticipate, based on their sequence diversity, that capped peptides will engage a diverse set of downstream receptor molecules to regulate pleotropic aspects of cell-cell communication and mammalian physiology.

## Acknowledgements

We thank members of the Long and Svensson labs for helpful discussions. This work was supported by the NIH (DK124265 and DK130541 to JZL; DK125260, DK11916, and DK116074 to KJS; GM113854 to VLL), the Ono Pharma Foundation (research grant to JZL), and the Stanford Wu Tsai Human Performance Alliance (research grant to JZL).

## Author contributions

Conceptualization (ALW, KJS, JZL); Methodology and software (ALW); Investigation (ALW, HZA, LV, VLL, JT, WW); Writing – Original Draft (ALW, JZL); Writing – Reviewing & Editing (ALW, HZA, LV, VLL, JT, WW, KJS, JZL), Supervision (JZL), Funding Acquisition (VLL, KJS, JZL).

## Methods

### Resource availability

#### Lead contact

Further information and requests for resources and reagents should be directed to and will be fulfilled by the lead contact, Jonathan Long.

#### Materials availability statement

Peptides generated in this study can be directly requested by email to the lead contact.

#### Data and code availability

Code for capped peptide prediction was deposited to GitHub (https://github.com/amandawigg/Capped-Peptides). All raw mass spectrometry data in Agilent .d file format were deposited to Mendeley Data (https://data.mendeley.com/v1/datasets/rcm9k9d2by/draft?preview=1).

### Experimental model and subject details

#### Mice and treatments

Animal experiments were performed according to a procedure approved by the Stanford University Administrative Panel on Laboratory Animal Care. Mice were maintained in 12-h light-dark cycles at 22°C and about 50% relative humidity and fed a standard irradiated rodent diet. Where indicated, a high-fat diet (D12492, Research Diets 60% kcal from fat) was used. Male C57BL/6J (stock number 000664) and male C57BL/6J DIO mice were purchased from the Jackson Laboratory (stock number 380050). For studies in high fat diet-fed mice, peptides were dissolved in 18:1:1 (by volume) of saline:Kolliphor EL (Sigma Aldrich):DMSO and administered to mice by intraperitoneal injections at a volume of 10 µl/g at the indicated doses for the indicated times. For lipopolysacchardide injection, LPS (Sigma, #L2880-10MG) was dissolved in saline and administered to mice at a volume of 5 µl/g at indicated dose. For fasting, food was removed from mice for 16 h. For running, a six-lane Columbus Instruments animal treadmill (product 1055-SRM-D65) was used with following 1 h protocol: 10 min at 6 m/min, 50 min at 18 m/min, and increase every 2 min by 2 m/min for the last 10 minutes, all at 12° incline. For all treatment experiments, mice were mock injected with the vehicle for 3-5 days until body weights were stabilized. Heparin plasm was harvested by submandibular bleed. For all experiments, mice were randomly assigned to treatment groups. Experimenters were not blinded to groups.

### Method details

#### Uniprot dataset curation

Lists of classically secreted proteins was obtained from Uniprot using the keyword “secreted” and filtering for either human or mouse species. Known peptide hormone sequences were obtained from Uniprot by first filtering for proteins annotated with keyword “hormone” for function and subsequently extracting out the specific hormone sequences from the peptides listed under the PTM annotations.

#### Computational prediction of capped peptides

Capped peptide prediction was accomplished using an in-house custom algorithm written in python (see code availability section). First, a list of classically secreted proteins was obtained from Uniprot filtering for species and the keyword “secreted.” Next, C-terminal amidation motifs were identified based on a GKR or GRR sequence indicative of dibasic cleavage and amidation upstream of the glycine. N-terminal pyroglutamylation was identified by searching for Q residues within 20 amino acids upstream of the amidation motif, and capped peptides were predicted to be the inclusive sequence between the N-terminal (pyro)glutamine and the C-terminal amidation.

#### BioGPS and GTExexpression analysis

For mouse, raw expression levels of precursor genes were obtained from the BioGPS dataset, GeneAtlas MOE430, gcrma. Replicates for the same tissue or cell type were averaged, and the relative expression was generated by normalizing the total of all tissue expression for a gene to 1. Next, the log of the relative expression was taken. An h-clustered heat map was made with the heatmap.2 function in gplots package in R using the z-score of the log(relative expression). For human, median gene-level expression was obtained from Gtex Analysis V8 and the heat map with h-clustered z-scores were similarly generated with heatmap.2 function in R.

#### Solid-phase synthesis of capped peptides

Capped peptides (human and mouse) were custom synthesized by Fmoc solid phase synthesis with Rink amide resin, or Wang resin for carboxy terminal CAP-TAC1 (Elim Biopharm, Hayward, CA). The crude product was used for all synthetic standards and validated by mass spectrometry. For functional studies, peptides were purified by HPLC to >90% purity. Identity and purity were verified by MS spectra and HPLC trace (Quality Control by Elim Biopharm, Hayward, CA, see SI).

#### Plasma and authentic peptide standards preparation for peptidomics

Plasma peptidomics preparations were adapted from (Ma et al., 2016). 1 µl of protease inhibitor (HALT, ThermoFisher, #78429) was added per 100 µl of plasma. Plasma was diluted 1:6 plasma:Tris-HCl buffer (100 mM Tris-HCl, pH 8.2) and boiled at 95°C for 10 minutes. In total, 1 ml of pooled plasma was used per replicate. 1 mM dithiothreitol (DTT, ThermoFisher, #FERR0861) was added, samples were vortexed and incubated for 50 minutes at 60°C. Iodoacetamide (IA, Sigma #I6125-5G) was added for a final concentration of 5 mM and incubated at room temperature for 1 hour in the dark. Formic acid (>95%, Sigma, #F0507-500ML) was added to 0.2% final concentration. Samples were centrifuged at 15,000 rpm for 20 min. Supernatants were concentrated with C8 columns (Waters, WAT054965), washed/desalted with water, and eluted in 100 µl of 80% acetonitrile. Samples were centrifuged at 15,000 rpm for 10 min. Supernatant was collected for liquid chromatography-mass spectrometry (LC-MS) analysis. All authentic peptide standards were pooled into 1 ml of Tris-HCl buffer (100 mM Tris-HCl, pH 8.2) and prepared in the same way as described above.

#### LC-MS detection of capped peptides

LC-MS was performed on an Agilent 6520 Quadrupole time-of-flight LC-MS instrument. MS analysis was performed using electrospray ionization (ESI) in positive mode. The dual ESI source parameters were set as follows: the gas temperature at 325°C, the drying gas flow rate at 13 l/min, the nebulizer pressure at 30 psig, the capillary voltage at 4,000 V, and the fragmentor voltage at 175 V. Separation of peptides was conducted using a C18 column (Agilent, #959961-902) with reverse phase chromatography. Mobile phases were as follows: buffer A, 100% water with 0.1% formic acid; buffer B, 90:10 acetonitrile:water with 0.1% formic acid. The LC gradient started at 95% A with a flow rate of 0.7 ml/min from 0 to 3 minutes. The gradient was then linearly increased to 60%A/40%B from 3 to 28 minutes and subsequently flushed at 5%A/95%B for 4 minutes and equilibrated back to 95%A/5%B for 6 minutes all at a flow rate of 0.7 ml/min. A positive capped peptide detection required a peak of exact mass (± 50 ppm) with total integration area of >1000 ion count and elution at the same retention time as the corresponding authentic synthetic standard. Exact masses and retention times of all detected peptides are listed in **Tables S2 and S5**. Quantification of each peptide was done by manual integration (AUC) in Agilent Mass Hunter software and compared to a standard curve to estimate plasma concentration.

#### Targeted LC-MS/MS

Targeted LC-MS/MS spectra were obtained using Agilent 6545 Quadrupole time-of-flight LC-MS instrument. The dual ESI source parameters were set as follows: the gas temperature at 325°C, the drying gas flow rate at 13 l/min, the nebulizer pressure at 30 psig, the capillary voltage at 4,000 V, the fragmentor voltage at 185 V, the sheath gas temperature at 350C, and the sheath gas flow rate at 11 l/min. The LC separation was done as described above. For CAP-GDF15, targeted LC-MS/MS method fragmented the +2 ion (m/z = 682.4) over the entire run with a collision energy of 40 and a narrow isolation width (∼1.3 m/z). For CAP-TAC1, targeted LC-MS/MS method fragmented the +1 ion (m/z = 724.3) over the entire run with a collision energy of 25 and a narrow isolation width (∼1.3 m/z).

#### BlastP alignment of capped peptides with whole mouse proteome

All detected mouse capped peptides were BLASTed to the mouse proteome (UniprotKB/Swiss-prot, taxid: 10090), using the BLOSUM62 matrix. 100% alignment was determined if the identities are 100%, positives are 100%, and gaps are 0%.

#### TACR agonist assay

Dose-response curves for CAP-TAC1 and positive control Substance P on the agonism of human TACR1/2/3 was measured by a Eurofins Discovery using human TACR1/2/3-transfected PathHunter beta-arrestin CHO-K1 cells.

#### Half-life calculations of CAP-TAC1 in plasma

10 µM of either synthetic CAP-TAC1 or SP (Sigma, #S6883) was incubated with 100 µl of mouse plasma. Samples were incubated at 37°C for 0, 10, 30, 60, or 90 minutes (N=3 for each time point). At indicated time point, 1 µl of HALT protease inhibitor was immediately added, and plasma was boiled and prepared using the peptidomics workflow described above. LC-MS spectra were obtained using Agilent 6545 Quadrupole time-of-flight LC-MS instrument as described above. Relative peptide levels were determined by total ion count area of exact mass (± 50 ppm, m/z = 724.3, +1 for CAP-TAC1 and m/z = 674.4, +1 for Substance P) peak that coeluted with synthetic standards. Basal plasma concentrations of Substance P and CAP-TAC1 were determined in samples with no synthetic peptide spiked in (N=3) with a standard curve comparison. Half-lives were calculated using an exponential decay fit.

#### Metabolic chamber studies

For acute energy expenditure studies, food intake, RER, movement, VO2, and VCO2 were collected with CLAMS Oxymax Metabolic Cages. Mice were placed individual metabolic cages for 24 hours prior to experiment. Mice were injected with either vehicle, 50 mg/kg CAP-GDF15, or 50 mg/kg scrambled CAP-GDF15 at 5 pm at T=0. Food intake, RER, movement, VO2, and VCO2 were collected every 7 minutes for 16 hours immediately after injection. All experiments were done with 3-5 month old DIO mice (Jackson, stock 380050), fed high fat diet ad libitum.

#### Sequence alignment for capped peptides

Sequence alignment was performed with EMBL-EBI Clustal Omega tool using the guided phylotree output (Madeira et al., 2022).

### Quantification and statistical analysis

Statistical analysis was performed in Prism 9.3.1. Student’s t test was used for pair-wise comparisons, and ANOVA was used for time course energy expenditure experiments were noted. Statistical significance was set at P<0.05. The specific test, P value symbol and error bar meaning, definition of center, and number of replicates are noted in figure legends.

## Supplemental Information

**Fig. S1.**
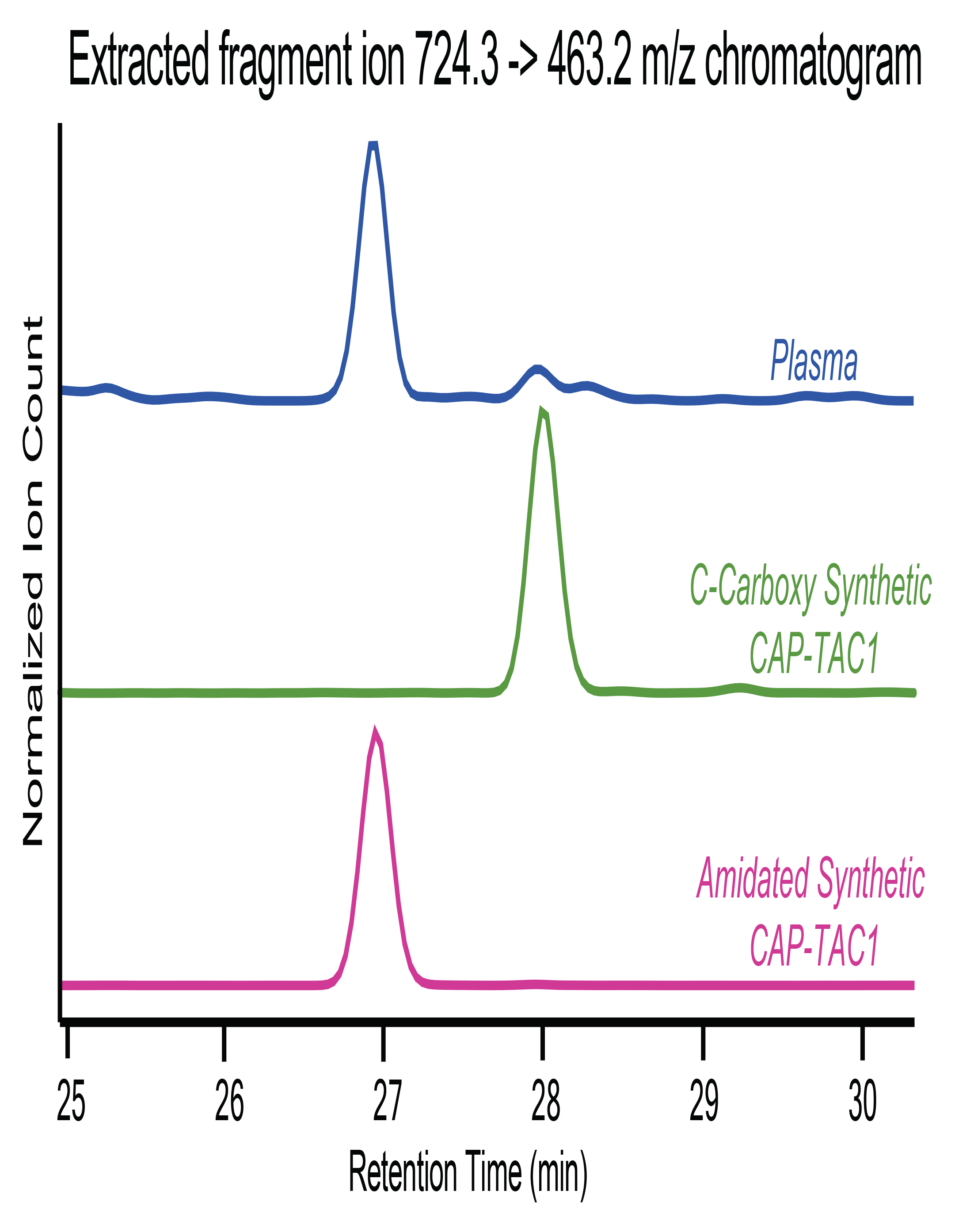
Comparison of C-terminal amidated and carboxy CAP-TAC1 with targeted LCMSMS method. The 463.2 m/z b4 extracted ion chromatograms for mouse plasma, the carboxy synthetic standard, and amidated synthetic standard (m/z = 724.3 parent ion with 1.3 m/z isolation window).

**Fig. S2.**
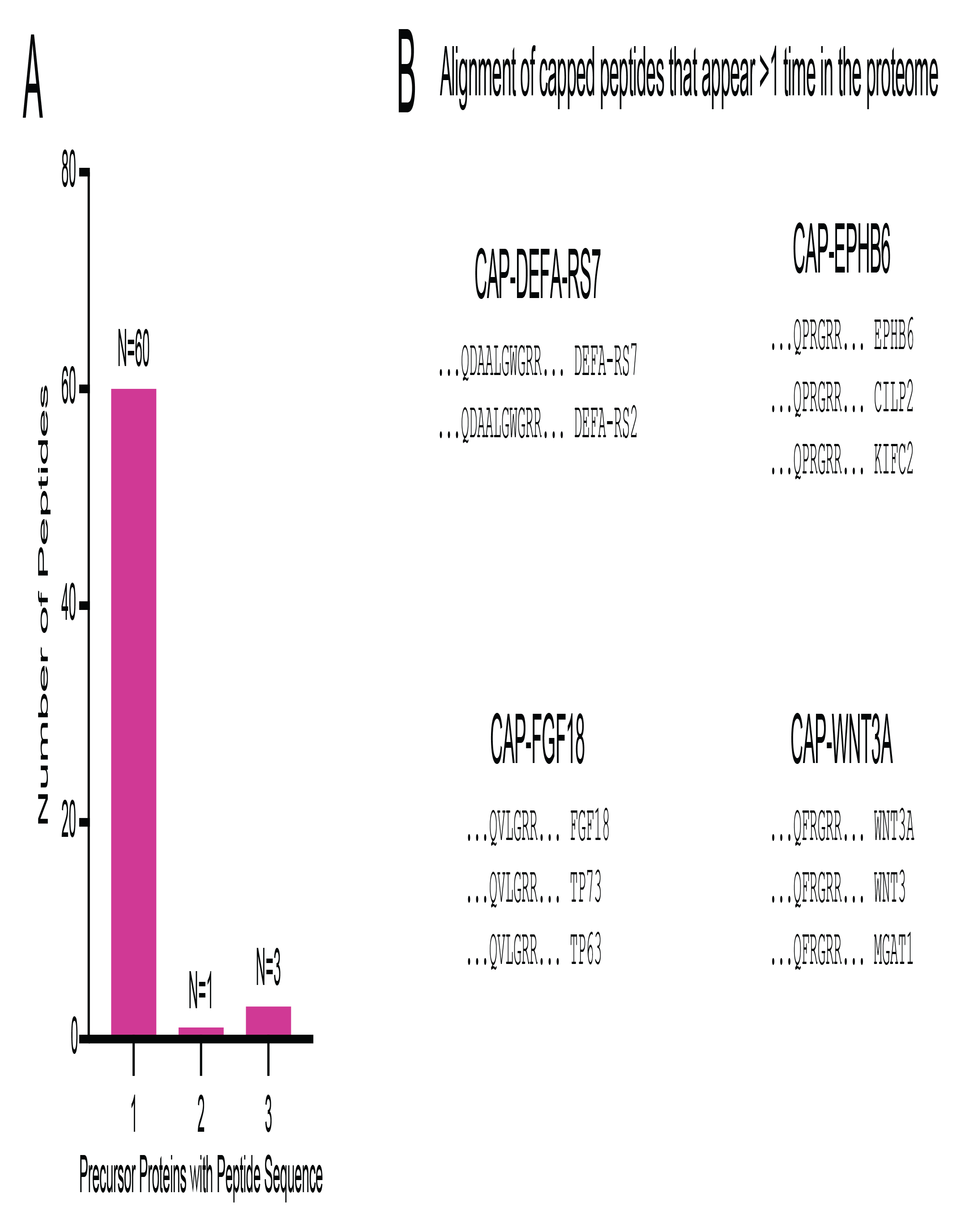
Determination of capped peptide sequence repeats in whole mouse proteome. Distribution of the number of precursor proteins (1, 2, or 3) the detected mouse capped peptide sequences (detected sequence + glycine-dibasic motif, N=64) are found in by a BlastP search.

**Fig. S3.**
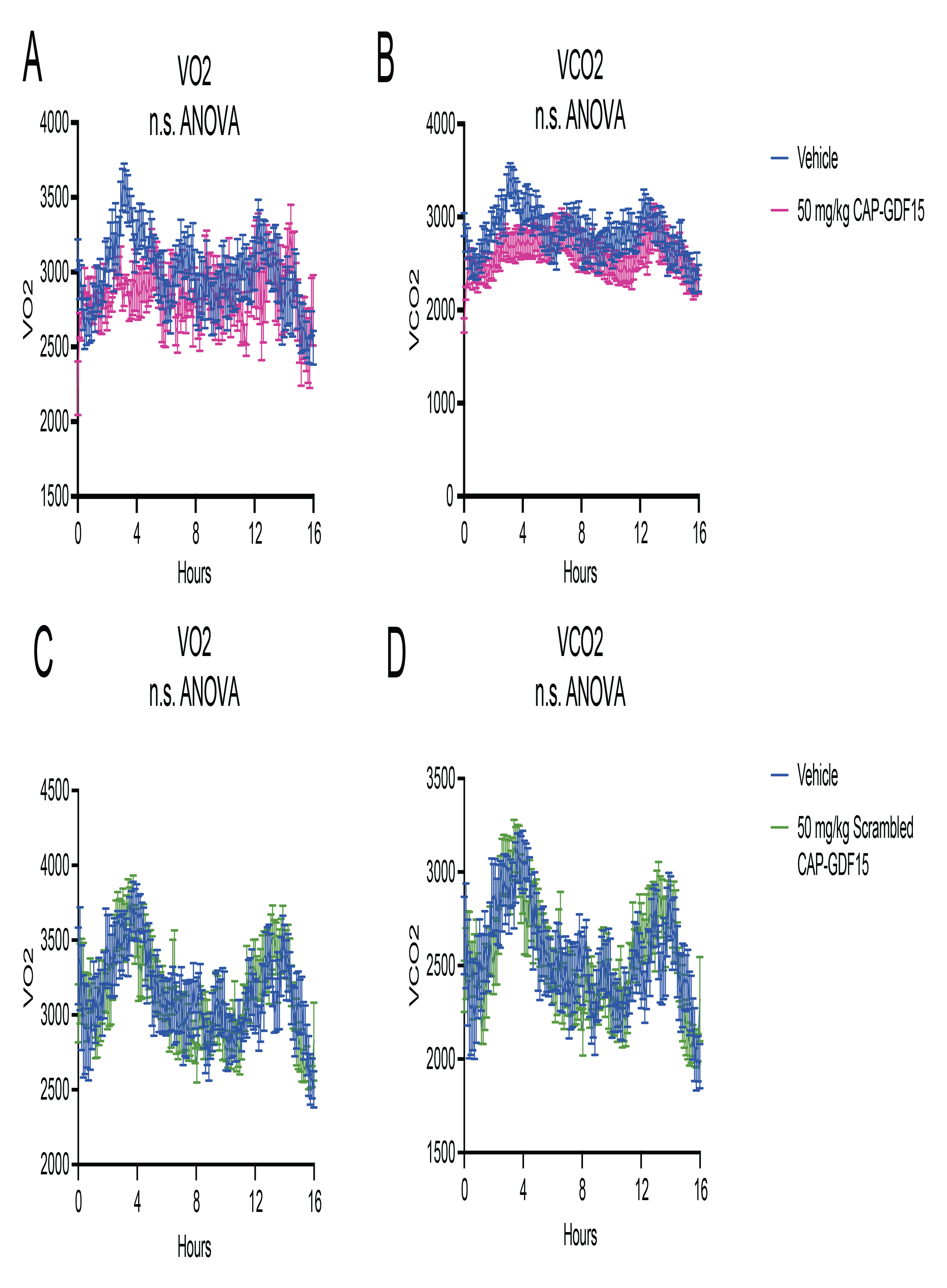
Additional measurements from metabolic chambers of mice treated with CAP-GDF15. (A,B) VO2 (A) and VCO2 (B) of 12-16-week diet-induced obese male mice following a single treatment of CAP-GDF15 (50 mg/kg, intraperitoneal) or vehicle control. (C,D) VO2 (C) and VCO2 (D) of 12-16-week diet-induced obese male mice following a single treatment of scrambled CAP-GDF15 (50 mg/kg, intraperitoneal, scrambled sequence: pQGLEALRARLRV-NH2) or vehicle control. Data are shown as means ± SEM. For (A-B), N=12/group; for (C-D), N=8/group. For (A-D), injection occurred at time T=0 (5:00pm) and data was collected for the subsequent 16 hrs.

**Fig. S4.**
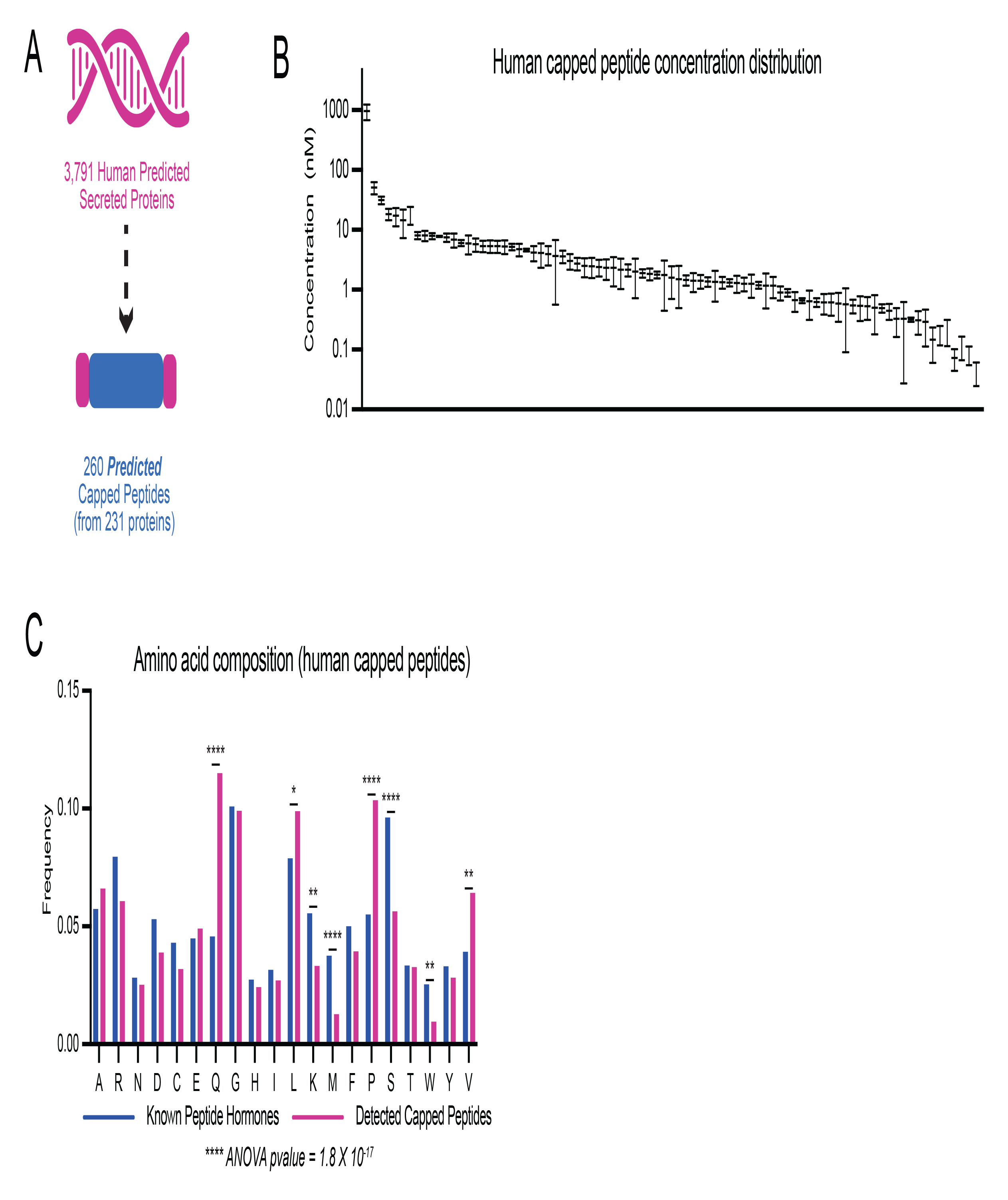
Prediction, detection, and composition analysis of human capped peptides. (A) Schematic of numbers of predicted human capped peptides. (B) Quantification of detectable human capped peptide concentrations in human plasma (N=3). (C) Comparison of frequency of each amino acid between human capped peptides and known peptide hormones. For (B), data are shown as means ± SEM, N=3.

**Fig. S5.**
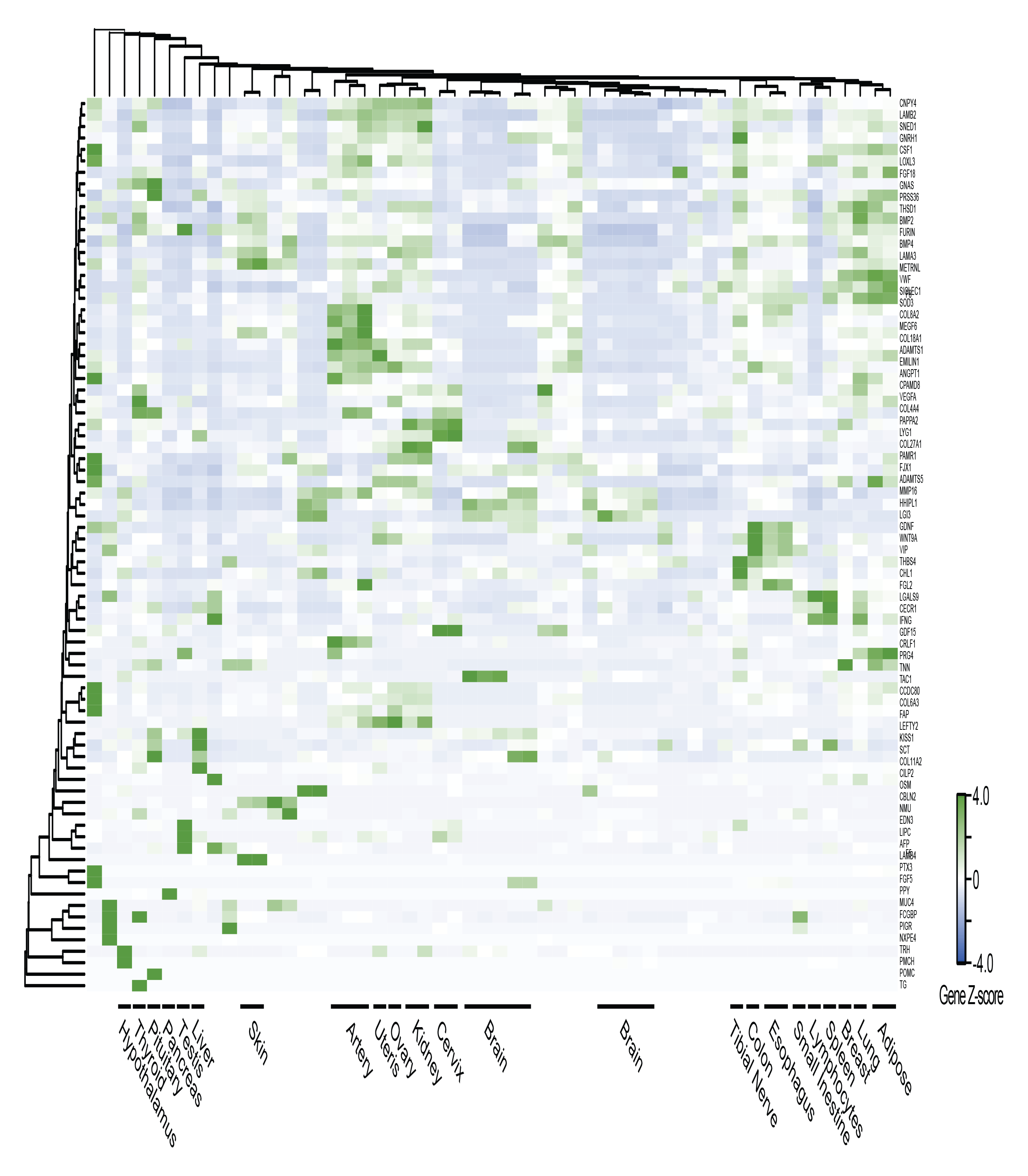
Tissue distribution of mRNAs for home genes corresponding to human capped peptide. H-clustered heat map of mRNA expression for capped peptide preproprecursor home genes across human tissues and cell types, using GTEx as the reference database.

**Table S1.** List of classically secreted mouse proteins from Uniprot.

**Table S2.** Predicted and detected mouse capped peptides.

**Table S3.** List of known peptide hormones and neuropeptides from Uniprot.

**Table S4.** List of classically secreted human proteins from Uniprot.

**Table S5.** Predicted and detected human capped peptides.

